# Dual agonist and antagonist muscle vibration produces a bias in end point with no change in variability

**DOI:** 10.1101/2025.03.03.641240

**Authors:** Gregg Eschelmuller, J. Timothy Inglis, Hyosub Kim, Romeo Chua

**Affiliations:** School of Kinesiology, University of British Columbia, Vancouver, British Columbia, Canada; Graduate Program in Neuroscience, University of British Columbia, Vancouver, British Columbia, Canada

## Abstract

Muscle spindles provide critical proprioceptive feedback about muscle length to the central nervous system (CNS). Single muscle tendon vibration can stimulate muscle spindles, causing participants to misjudge limb position, while dual muscle tendon vibration is thought to produce a noisy proprioceptive system. It is currently unclear exactly how the CNS uses kinesthetic feedback from the agonist and antagonist muscles during target-directed reaches. The purpose of the current project was to investigate the effects of agonist, antagonist, and dual agonist/antagonist during target-directed reaching. Using an elbow extension task, we found that antagonist muscle vibration produced an undershooting effect relative to the no-vibration control, while agonist muscle vibration produced an overshooting effect relative to the no-vibration control. Neither of the single muscle vibrations produced any change in the variable error of the movements. While it was originally hypothesized that dual agonist/antagonist vibration would increase participants variable error with no change in bias, the opposite was found. Participants undershot relative to the no-vibration control with no change in variable error. Overall, the results from this study suggest that dual vibration does not necessarily create a noisy proprioceptive system but can produce a bias in end point.

## Introduction

Proprioceptive feedback from mechanoreceptors in our muscles is critical to our ability to move our bodies through space. While there are multiple receptors that contribute to our proprioceptive senses (Proske and Gandevia 2012), the muscle spindles are thought to be our primary kinesthetic receptor (Proske and Gandevia 2018). Kinesthesia is strictly defined as our sense of movement, but it more commonly is used to describe both our sense of movement and position, which is the operational definition used here. One way to study the role of muscle spindle feedback in motor control is to artificially activate them with muscle tendon or muscle belly vibration (Burke et al. 1976a, b), which produces illusions of position and movement (Goodwin et al. 1972; Inglis and Frank 1990; Inglis et al. 1991; Cordo et al. 1995). Specifically, vibration of a muscle or muscle tendon will induce a kinesthetic illusion that the muscle is in a longer state than it actually is. For example, when participants are asked to make a target-directed movement, vibration of the antagonist (lengthening) muscle, produces an undershooting effect compared to the no-vibration control (Inglis and Frank 1990; Cordo et al. 1995; Eschelmuller et al. 2023). However, vibration of the agonist (shortening) muscle in this situation produces no kinesthetic illusion (Inglis and Frank 1990). This indicates that the estimate of joint angle is primarily computed based on the afferent input from the lengthening muscle, and not the shortening muscle. While α-γ coactivation (parallel activation and α and γ motoneurons), may be able to keep the muscle spindle taut during slow shortening contractions, this may not take up enough slack for the muscle spindle to continue to provide reliable afferent feedback, and therefore, the central nervous system (CNS) may favour the muscle spindle afferent input from the lengthening muscle.

Dual agonist/antagonist vibration has been suggested to degrade proprioception in a variety of proprioceptive tasks, including joint angle matching at the wrist (Bock et al. 2007). Bock et al. (2007) demonstrated that, when performing angle matching with and without dual agonist/antagonist vibration, the absolute error increased in the vibration condition. This degradation of performance is presumably due to a “busy line” effect in the muscle spindles, rendering the afferent signal noisier (Roll et al. 1989). Since the muscle spindles would be firing in sync with the tendon vibration, their ability to encode the actual movements of the muscle would decrease. However, a few key questions remain about the use of dual tendon vibration to degrade proprioception. First, it is unclear how dual agonist/antagonist vibration would affect performance in a target-directed reaching task, where the feedback must be utilized by the CNS to complete a motor action. Additionally, it is unclear based on Bock et al., (2007), if dual agonist/antagonist tendon vibration induced a change in the constant or the variable error. Previous work has demonstrated that single muscle vibration will typically demonstrate a change in the constant error, but no change in the variable error (Inglis and Frank 1990; Eschelmuller et al. 2023). If the dual vibration degrades proprioception we would expect to see changes in the variable error, with no change in constant error as previous work has demonstrated that there are little to no perceptual illusions with dual vibration at the same frequencies, in an angle matching task (Gilhodes et al. 1986). Therefore, it may be that dual vibration produces a change in the variable error and not the constant error in proprioceptive tasks. However, if the CNS only uses proprioceptive feedback from the lengthening muscle to estimate joint angle when performing a targeting task (Inglis and Frank 1990), when we vibrate both tendons, it is unclear what will happen when one muscle is shortening, and one is lengthening. If the CNS is only using the afferent information from the lengthening muscle to compute joint angle, then the dual vibration should produce similar effects as single muscle vibration of the lengthening muscle, as the vibration-induced feedback from the agonist would not be used in the joint angle estimate. Therefore, the purpose of the current study was to investigate the effects of agonist, antagonist, and dual agonist/antagonist tendon vibration in a target-directed task.

## Methods

### Participants

15 participants (age: 22.46 ± 2.63; 12 female) free of any neurological or musculoskeletal disorders were recruited to participate in this study. Procedures were approved through the University of British Columbia behavioural research ethics board. Informed written consent was obtained from all participants. Participants were remunerated $10 for their participation.

### Experimental Setup

Participants were seated in a chair that allowed for height and trunk angle adjustments. Participants were seated with their right arm on a manipulandum that allowed for flexion and extension of their elbow, but limited movement of the wrist and shoulder. The arm was pronated so the hand lay flat on a metal palm rest allowing participants to align their middle finger forward along the manipulandum handle. A potentiometer (Vishay Spectrol 157) was mounted in the manipulandum axis to record elbow angle in volts, which was then calibrated to degrees to record elbow angle. The height and angle of the chair was adjusted so that the arm was as close to horizontal as possible, without producing discomfort for the participant. To produce tendon vibration, two muscle vibrators, which consisted of an eccentric-mass rotary motor in a cylindrical plastic case measuring 25 × 50 mm, were secured on the right biceps brachii and triceps brachii tendons with a tensor bandage. The vibrators were driven at ∼90-95 Hz by a DC power supply (Zhaoxin PS-3005D), controlled through Python code via a NI USB-60001 multifunctional DAQ (National Instruments, USA). The muscle that each individual vibrator was secured to was swapped between participants to control for any subtle differences in vibration between different vibrators. All experimental stimuli were controlled using custom Python scripts using the Psychopy library (Peirce et al. 2019). Data were collected and streamed into Python using an NI USB-60001 multifunctional DAQ (National Instruments, USA), using the NI-DAQmx application programming interface. A curved (1000r) 27-inch 1080p monitor (MSI optix G271C, Micro-Star Int’l Co., Ltd) was placed in front of the participant and was used to display all experimental stimuli. The placement of the monitor was such that it was in line with the curvature of the manipulandum. A black board was secured over top of the arm to occlude vision of the arm and hand from the participant.

### Experimental Procedures

To investigate the effects of muscle tendon vibration, participants made elbow extension movements of their unseen right arm to a visual target, which consisted of two vertical red lines spanning the height of the monitor, spaced 1 cm apart. The goal of the participant was to align their middle finger between these two red lines. The starting elbow angle was ∼40 degrees of flexion, which was purposely jittered by the experimenter by ± ∼3 degrees (home zone). The target position had a mean angle of 100 degrees of flexion, with random jitter of ± ∼3 degrees. The jitter in the start position and target position was to avoid participants memorizing the movement distance. Participants were asked to stop their movement at the target and wait there, and the experimenter would return them to the home zone. On the return path the experimenter would move the participant’s arm through a few random cycles of flexion and extension to eliminate any cues of the end position. Participants first completed a practice block for 30 trials, where they received terminal feedback of their end point to ensure they understood and could complete the task with confidence. Following this practice period, participants completed a baseline block that had 30 trials with no terminal feedback. Next, participants completed the main experimental block, where they completed 6 mini blocks that consisted of 10 trials with no vibration and terminal feedback, followed by 30 trials with no terminal feedback and randomly delivered tendon vibration. Within these no-feedback trials, 15 had no vibration and 15 had tendon vibration, which was subdivided into agonist vibration (triceps), antagonist vibration (biceps), and dual agonist/antagonist vibration. Trials were randomized within each mini block. The feedback blocks in the mini blocks were to ensure participants remained confident and accurate in their movements, and the no-feedback blocks were the primary blocks to evaluate the effects of tendon vibration. Following the main experimental block, participants performed a post block, which was the same as the baseline (30 trials no feedback, no vibration). When the participants received terminal feedback, it was presented at the end of the movement (velocity below ∼1 deg/sec) as a stationary solid green line and stayed on the screen for 500 ms.

During each trial, participants would see a yellow circle in the middle of the screen which turned green once the experimenter moved them into the home zone. After a random delay (1000 ms to 2000 ms), the target would appear, and participants were asked to make one smooth movement and stop at the target. The emphasis was to produce a single movement without adjustment. The participants practiced the movements and were given verbal feedback regarding their velocity to keep it between 40-60 deg/second. During the experimental trials no feedback was given to avoid influencing the effects of vibration. During vibration trials, the vibrator turned on at target onset and stayed on until the end of movement. The end point was calculated as the point when movement velocity dropped below ∼1 deg/second.

### Data Analysis

All data analysis and visualization were done in Python and used the following libraries: pandas (The pandas development team, 2024), Numpy (Harris et al. 2020), Matplotlib (Hunter 2007), and Seaborn (Waskom 2021). Trials were removed if the mean velocity was less than 20 degrees/second or higher than 80 degrees/second. This led to ∼1.8% of trials being removed on average (range: 0 - 7.9%). Next, an error score was calculated between the end point and the target location. The convention of target position subtracted from the end point was used, which means that a negative value indicates that the participant undershot the target. This error will be referred to as ‘target error’. To calculate the effects of tendon vibration, end point data were first averaged across each condition to produce mean values for each participant, then the no-vibration mean was subtracted from each vibration condition for each participant separately. This error will be referred to as ‘vibration error’. Again, a negative value indicates that the vibration condition produced undershooting relative to the no-vibration condition. To investigate if exposure to the experimental block affected participants’ performance, the baseline and post blocks that had no vibration and no feedback were compared. Variable error was calculated as the standard deviation of the target errors. Average velocity was calculated as the movement distance divided by movement time for each trial separately.

### Statistical analysis

Statistical analysis was done in the JASP statistical software (JASP Team 2024). Target error, variable error, and mean velocity were analyzed using separate one-way repeated measures ANOVAs with the sole factor of vibration condition (biceps, triceps, dual, and no vibration). Significant main effects were followed up with Bonferroni-corrected paired t-tests. The pre and post no-vibration comparison was analyzed using a paired t-test. Statistical significance was set at p < 0.05 for all measurements and a Greenhouse–Geisser adjustment was applied when sphericity was violated. Uncorrected degrees of freedom are reported in the text.

## Results

In the no-vibration condition, participants were able to complete the task with a slight undershooting of the target by ∼2.3 degrees, and a variable error of ∼4 degrees. We trained participants at the beginning to move with a mean velocity in the range of 40-60 degrees/second, and on average the mean velocity was 49.8 degrees/second. Therefore, at baseline (no vibration), participants were able to complete the task with relatively good accuracy and movement characteristics. When looking at our control and vibration data, there was a significant effect of vibration condition on target error [F (3, 42) = 43.213, p < 0.001, 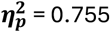] (Figure 2) but not on variable error [F (3, 42) = 0.762, p = 0.404, 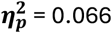] (Figure 3). When examining the target error, follow-up post-hoc tests revealed that biceps vibration resulted in a 4.22 (95% CI [2.37, 6.08]) degree undershoot relative to control (p < 0.001), while triceps vibration resulted in a 1.80 (95% CI [0.43, 3.17]) degree overshoot relative to control (p = 0.008). The dual vibration resulted in an undershoot of 1.67 (95% CI [0.20, 3.15]) degrees relative to control (p = 0.022). When comparing across vibration conditions, all pairwise comparisons were significantly different from each other (p < 0.05). See Error bars represent bootstrapped (10,000x) 95% confidence intervals.

**Figure 1.**
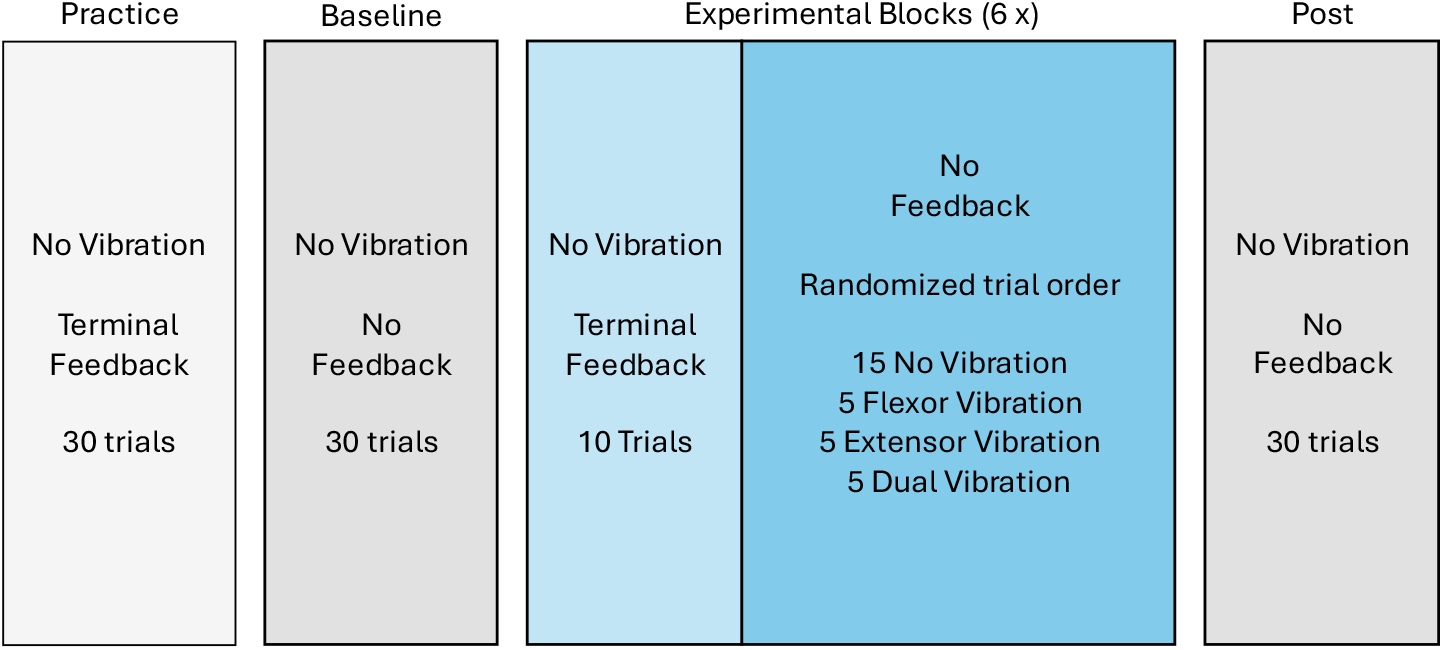
Experimental protocols. Note the experimental block was repeated 6 times.

**Figure 2.**
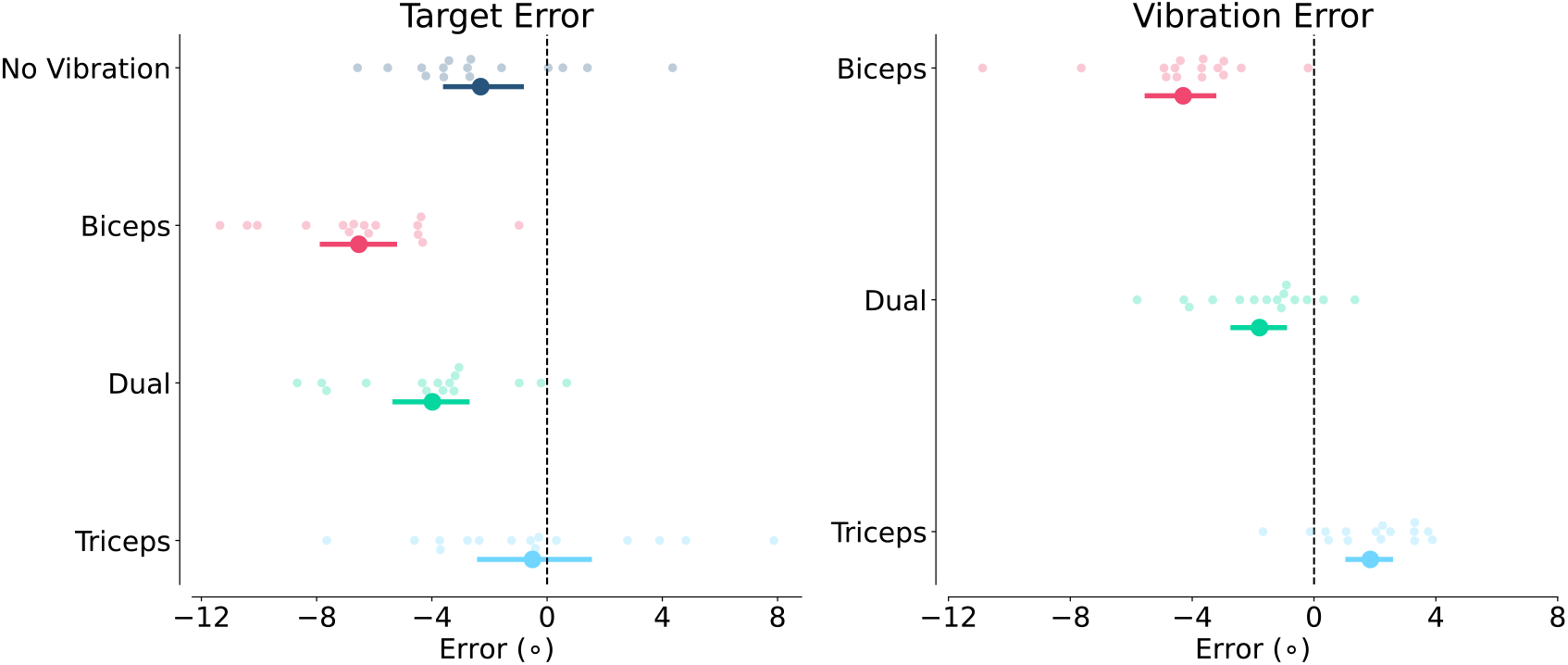
Mean target error (left), mean vibration-induced error (right). A negative target error indicates an undershooting of the target. By convention vibration error is calculated as no-vibration subtracted from vibration, which means a negative value is an undershooting relative to the no-vibration mean. Error bars represent bootstrapped (10,000x) 95% confidence intervals.

**Figure 3:**
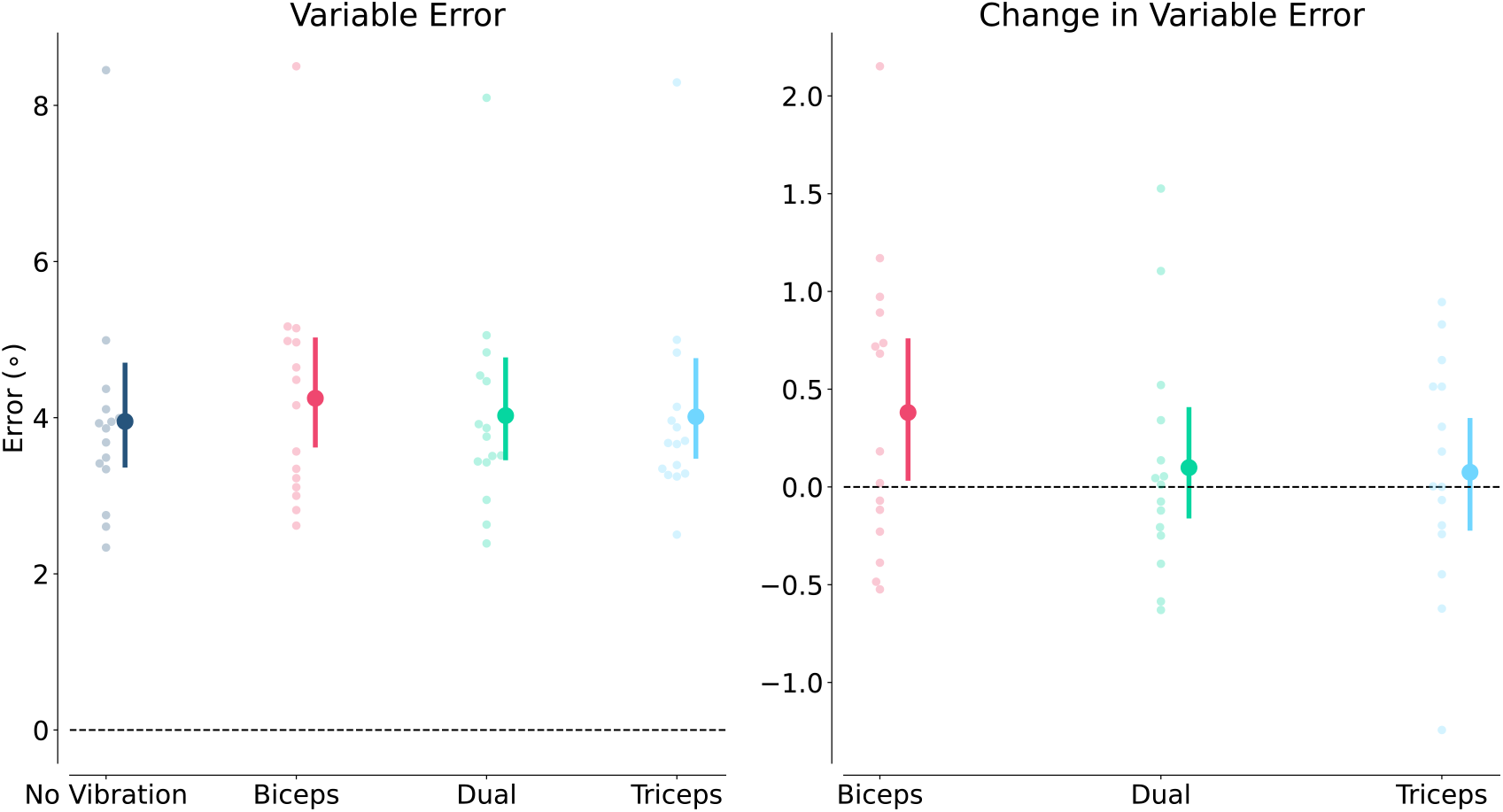
Mean target variable error (left), mean change in variable error (right). By convention change in variable error is calculated as no-vibration subtracted from vibration, which means a positive value is higher variability. Error bars represent bootstrapped (10,000x) 95% confidence intervals.

Table 1 for a full summary of the post-hoc analysis on target error.

**Table 1:**
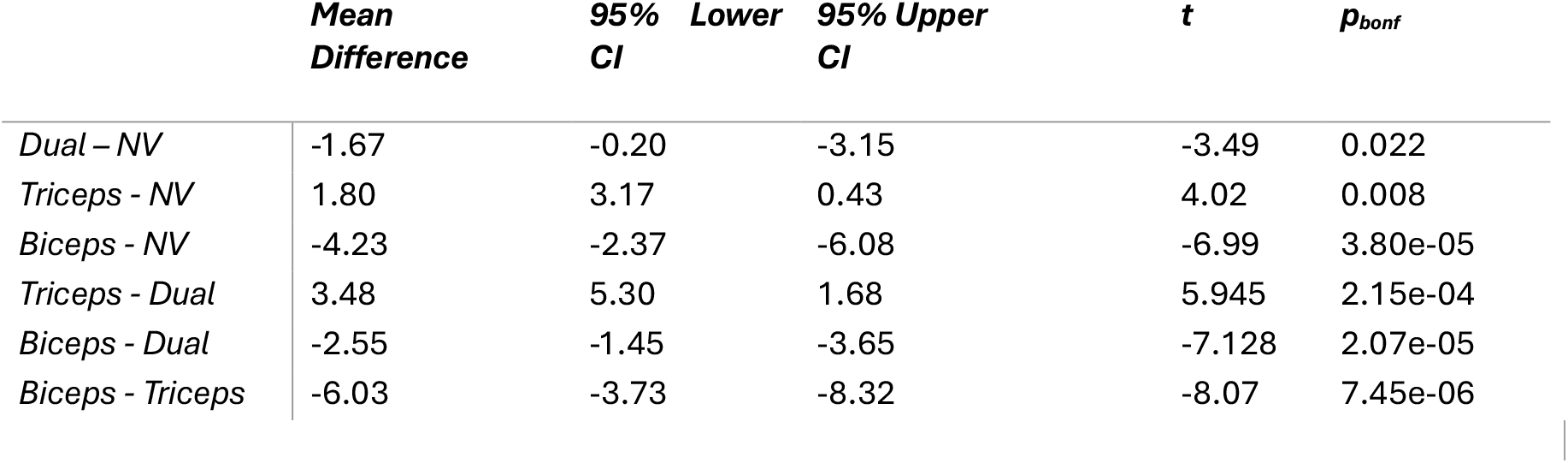
Post-hoc analysis of vibration-induced errors. Note. P-values and confidence intervals are adjusted for comparing a family of 6 estimates (Bonferroni method).

As vibration can produce changes in both end point and movement speed, we examined the change in mean velocity across vibration conditions. We found that vibration had a significant effect on mean velocity [F (3, 42) = 17.029, p < 0.001, 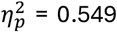] (Figure 4). Most notable, the biceps vibration condition produced slower movements compared to the no-vibration (p < 0.001), the triceps vibration (p < 0.001), and dual vibration conditions (p = 0.031). The dual vibration also resulted in slower movements compared to the no-vibration (p = 0.003) and the triceps vibration conditions (p = 0.005). There were no differences between the triceps vibration and the no-vibration condition (p = 1.00).

**Figure 4.**
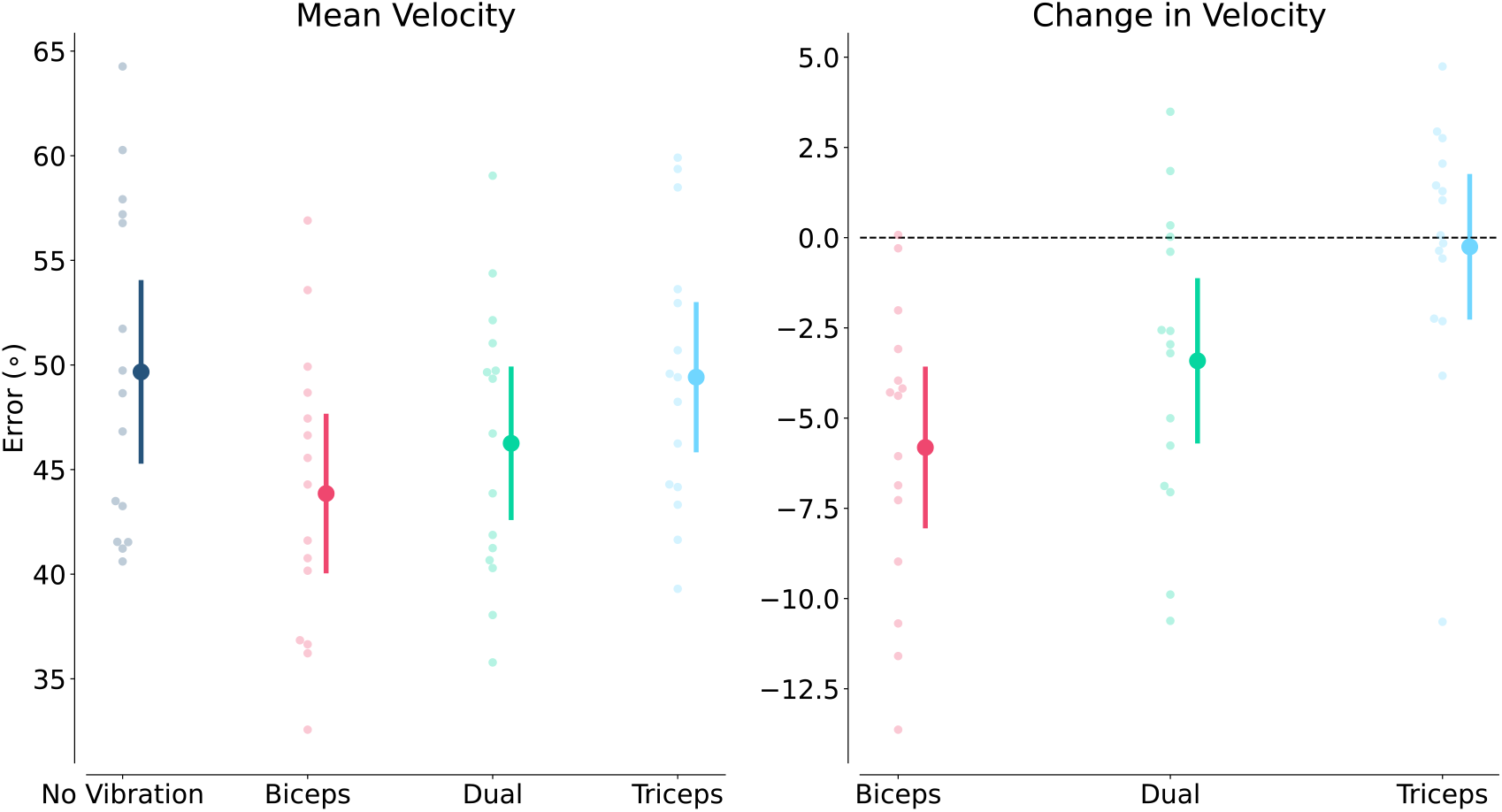
Mean velocity of movements (left) and change in velocity with vibration (right). By convention change in velocity is calculated as no-vibration subtracted from vibration, which means a negative value is a lower velocity. Error bars represent bootstrapped (10,000x) 95% confidence intervals.

When examining the performance in our baseline block and our post block, there were no significant differences in mean target error between blocks (p = 0.110) (Figure 5). This would indicate that the exposure to the vibration trials did not systematically affect overall performance when participants were tested in the no-feedback, no-vibration post block.

**Figure 5.**
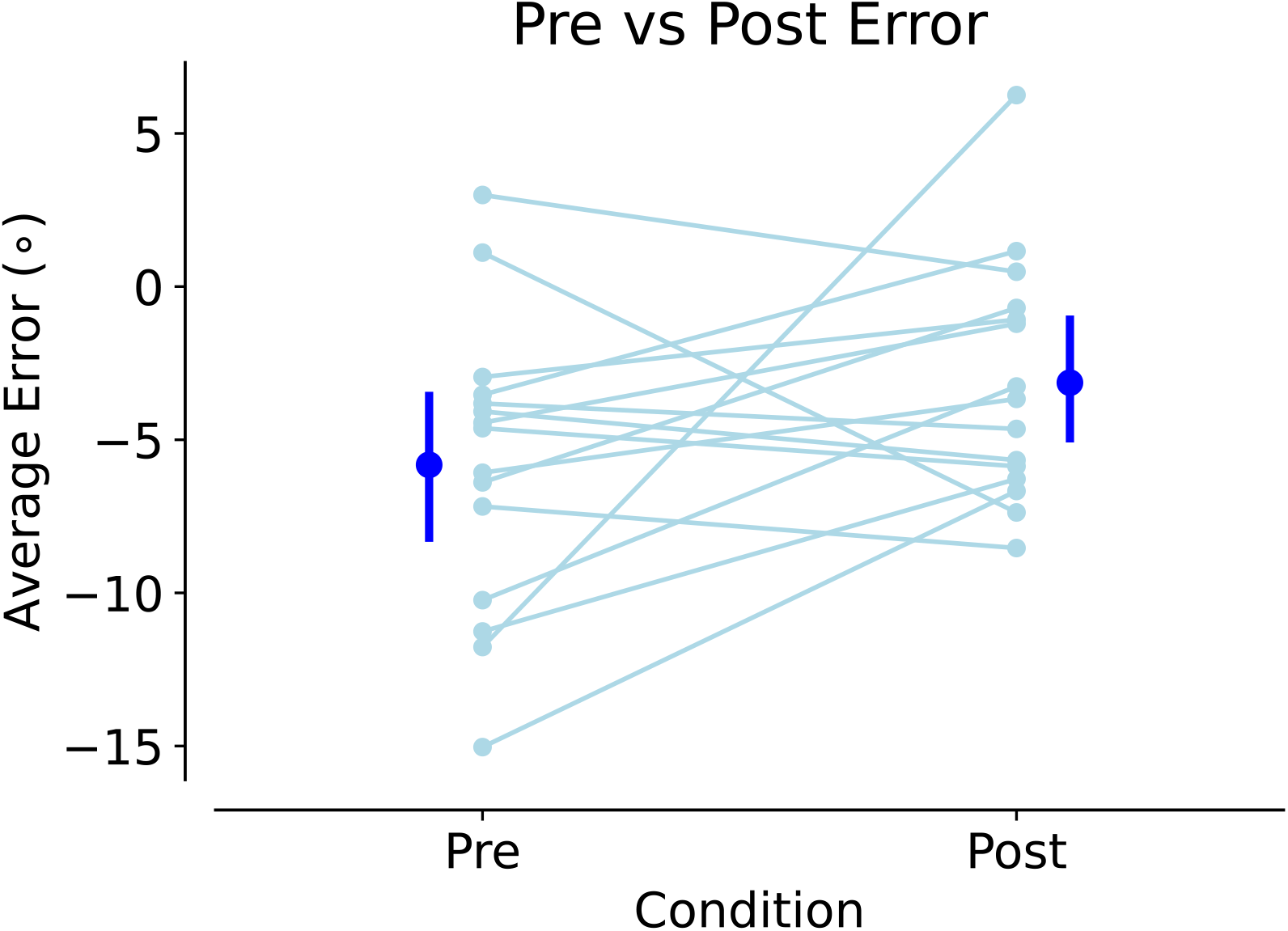
Pre and post vibration exposure errors. These data represent no-feedback with no-vibration target errors. There was no statistical difference between the pre and post errors.

To examine more closely the effects of dual vibration, the sum of the biceps and triceps vibration-induced errors were compared to the dual vibration error for each participant. When the sum of the biceps and triceps errors was compared to the dual vibration errors, there was a significant positive correlation between the two [r = 0.776 95% CI [0.44, 0.92]; p < 0.001]. Figure 6 shows the scatter plot of the dual vibration error against the triceps + biceps vibration errors, relative to the line of unity. If the effect of dual vibration is simply the algebraic sum of the individual effects of triceps and biceps vibration, the data points would fall along this line. While the data points generally follow the line of unity, the slope was less than 1, which could indicate a regression to the mean effect.

**Figure 6.**
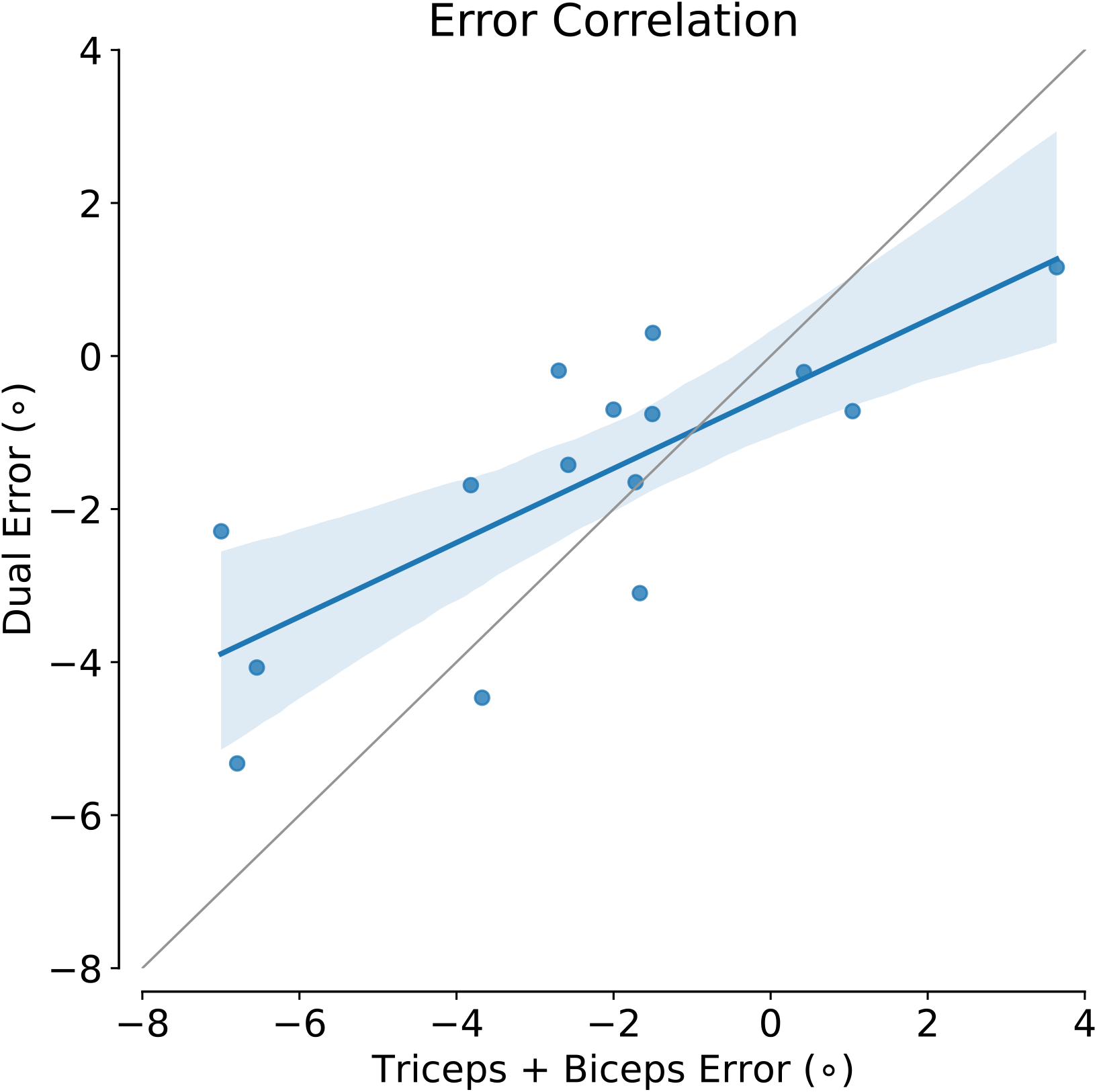
Dual vibration errors plotted against the sum of triceps and biceps vibration errors in degrees. Grey line represents unity line. If the dual error is equal to the sum of the triceps and biceps errors, data would fall along this line. r = 0.776 95% CI [0.438, 0.922]; p < 0.001

## Discussion

The objective of the current study was to investigate the effects of dual agonist/antagonist tendon vibration on performance in target-directed reaching. Using an elbow extension task, we were able to classify the effects of agonist, antagonist, and dual agonist/antagonist tendon vibration on both the constant error and the variable error. With single muscle vibration, the original hypotheses for this study were that 1) biceps brachii vibration would produce an undershooting effect relative to the control, and 2) the triceps vibration would produce no effect. Our results for the biceps vibration are in line with previous literature and our hypothesis that it would induce an undershooting effect relative to the no-vibration condition (negative vibration error). This result has been consistently demonstrated in the literature (i.e. Cordo et al., 1995; Inglis & Frank, 1990), and is proposed to be due to the vibration-induced increase in muscle spindle afferent activity, which leads to the perception of a longer muscle. Unlike previous results (Inglis and Frank 1990), our results indicate that vibration can induce an illusion in the shortening agonist muscle (triceps brachii). Specifically, in the triceps brachii vibration condition, participants tended to overshoot relative to the no-vibration condition. It was previously proposed that only the afferent information from the lengthening muscle was used by the CNS in targeting tasks (Inglis and Frank 1990); however, the data from this study indicate this is not the case. We suggest that when a muscle is shortening, the afferent code may become unreliable as the muscle spindles become unloaded and therefore provide an unreliable signal for joint angle. Potentially, the movements in this study were slow enough that the muscle spindles did not become fully unloaded, and therefore, the agonist was still able to provide some reliable information for the CNS. Previous work (Inglis and Frank 1990) using similar movement speeds found conflicting results, and therefore, it is unclear what is driving the discrepency in the results. Differences in other methodologies, such as vibration (model/make, amplitude), experimental task (bimanual vs unimanual), and experimental protocol (blocked vs randomized) could be causing the differences seen.

It has been previously suggested that dual agonist/antagonist vibration can ‘degrade’ proprioception (Bock et al. 2007); however, it was unclear whether this degradation was due to a bias in proprioception or if it was producing a “busy line” effect and making the information less reliable. We hypothesized that if it truly degraded proprioception, there should be a change in the variable error but no change in the constant error. The results from this study do not align with this idea, as we found that dual agonist/antagonist vibration produced no change in variable error but did cause participants to undershoot the target. When the data were further examined, we found that the errors within the dual vibration condition significantly correlated with the sum of the biceps and triceps errors for each participant. Previous work using an angle matching task has demonstrated that when the two muscles are vibrated at the same frequency, there is either no illusion or an illusion of very slow movement (Gilhodes et al. 1986). When the two vibrators were driven at different frequencies, illusions would reappear. It was argued that the perceptual illusions of the participants were due to the difference in vibration frequencies between the muscles. The present data are in line with this idea that the net effect of dual vibration is the sum of the individual vibration effects. In the current study, however, when both muscles were vibrated, participants still tended to undershoot the target. In the task that was used, participants extended the elbow to the target, and therefore, the muscle spindles in the biceps were lengthening and were likely very sensitive to vibration, while the muscle spindles in the triceps were shortening and were likely less sensitive to vibration (Burke et al. 1976a, b). Therefore, while the net effect of dual vibration is the weighted average of the individual vibrations, the overall behaviour still depends on the movement being performed. In this case, the illusions for biceps vibrations were larger in magnitude than the triceps vibrations, so their net sum was still that of undershooting. However, the slope of the data was slightly less than 1, indicating this simple model is not fully capturing the behaviour. In this paradigm, a slope of less than 1 would indicate that the magnitude of illusion in the dual vibration is slightly less than the sum of the biceps and triceps vibration. One potential explanation for this could be that when both muscles are vibrated, the vibration signals could be propagated to both agonists and antagonists leading to some destructive interference of the signals leading to less of the signal being detected by the muscle spindles. Regardless, the strong linear relationship between the sum of the biceps and triceps errors and the dual vibration errors indicate that the CNS is utilizing proprioceptive feedback from both agonist and antagonist muscles around a joint.

Neither the biceps nor triceps vibration conditions resulted in any change in variable error, indicating that both produced a kinesthetic bias without introducing any significant noise into their targeting performance, which is in line with the results from previous work (Inglis and Frank 1990; Eschelmuller et al. 2023). Interestingly, the dual vibration condition also did not induce an increase in variable error. It is unclear why the dual vibration did not increase variable error, as the vibration was likely reducing the ability of many of the muscle spindles in both muscles to encode the actual movement. Previous work has suggested that tendon vibration locks a sub population of muscle spindles to the vibration frequency (Burke et al. 1976b, a; Roll and Vedel 1982; Roll et al. 1989). However, if this were the case during dual vibration, a large population of the muscle spindles would not be able to encode the movements which would be expected to cause more variability in the movements. While there is a lack of data on muscle spindle recordings during larger goal-directed movements with vibration, it may be that instead of locking a subpopulation of muscle spindles, vibration increases the overall firing rates of the muscle spindles, but they are still able to encode the movement. In this case, the population code would indicate a slightly longer muscle length with vibration, but the spindles could still encode the relative muscle length changes. This theory would explain the consistent results of vibration inducing a bias with no change in variable error. However, it is important to note that while the muscle spindles are the primary kinesthetic receptor, cutaneous receptors around the joint along with the efferent prediction related to the motor command could also provide important information to determine joint angle (Proske and Gandevia 2018). It is possible that the dual vibration did induce noise into the muscle spindles, but the signals from the cutaneous receptors and the efferent prediction based on the motor command were able to compensate for this noise and therefore, no increase in variable error was present. In line with this idea, data in patients with hereditary sensory and autonomic neuropathy type III, who lack functional muscle spindles (Macefield et al. 2011), do not display deficits in angle matching tasks of the upper limb (Smith et al. 2020). The authors argue that this is because the cutaneous receptors around the joint were able to take on the role of the primary kinesthetic receptor in this task. The data from the current study cannot distinguish which mechanism is responsible for lack of change in variable error.

Lastly, we found that tendon vibration produced changes in participants’ movement velocities. Notably, in the biceps and dual vibration conditions, participants moved slower compared to the no-vibration and triceps vibration conditions. This is not an unexpected finding, as muscle spindles are thought to encode both static (muscle length) and dynamic (muscle length change) information (Proske and Gandevia 2012). In these two conditions, the activation of the lengthening muscle would produce the illusion of a more extended elbow position and also that the arm was moving into extension at a higher velocity, resulting in both an undershooting of final position and a slower movement. Interestingly, the triceps vibration condition did not produce an increase in movement velocity, even though it produced an overshooting effect of the target. From the current data, it is not possible to decompose a causal relationship between the movement velocities and end points in the biceps and dual vibration conditions as both the static and dynamic information provided by the muscle spindles is affected by muscle tendon vibration.

Overall, these findings provide important insight into how the CNS uses proprioceptive afferent information from both the agonists and antagonists during goal-directed reaching. The data suggest that the CNS uses all the information that is available from both the agonists and antagonists, but greater weight may be given to the lengthening antagonist muscle in targeting tasks as this feedback is more reliable. Dual vibration, while previously suggested to ‘degrade’ proprioception, seems to represent the net sum of the agonist/antagonist vibration, so the effects will depend on the task that is being performed. However, since none of the vibration conditions produced any change in the variable error, it is unclear whether it truly degrades proprioception or just introduces a bias. We are currently examining the relationship between the movement type and the effects of tendon vibration, to further understand how the CNS uses proprioceptive afferent feedback to control goal-directed movements.

## Acknowledgements

The authors thank Annika Szarka and Nick Butler for their help in data collection.

## Funding

This work was supported by the Natural Sciences and Engineering Research Council to JTI [Grant number: 2017-04504] and RC [Grant number: 2019-04513] and a graduate research award to GE.

## Data availability

The datasets generated during and/or analyzed during the current study are available from the corresponding author on reasonable request.

## Conflict of interest

The authors declare that they have no conflict of interest.

## Ethical approval and consent to participate

All procedures used in this study were approved by the behaviour research ethics board at the University of British Columbia. Freely given informed written consent was obtained from all participants prior to participation in any study procedures.

## Consent for publication

All participants consented for their de-identified data to be published.

## Notes

### Competing Interest Statement

The authors have declared no competing interest.

### Summary of Updates

There was a formatting issue in which Figure 3 appeared twice. The duplicate image has been removed.

